# Neural stem cells isolated from activated brain neurogenic niches after traumatic injury demonstrate distinct behaviors *in vitro*

**DOI:** 10.1101/2024.01.27.577567

**Authors:** Lars Erik Schiro, Ulrich Stefan Bauer, Christiana Bjorkli, Axel Sandvig, Ioanna Sandvig

## Abstract

The subventricular zone (SVZ) of the lateral ventricle and subgranular zone (SGZ) of the dentate gyrus (DG) in the hippocampus represents neurogenic niches within the brain housing distinct populations of neural stem cells (NSCs), known for their exclusive capacity to sustain neurogenesis in the adult mammalian CNS. These niches respond to traumatic brain injury (TBI) by becoming activated, leading to NSC proliferation, a small number of which subsequently migrate towards the injury site, and differentiate predominantly into astrocytes. Although the capacity of activated NSCs to differentiate into neurons appears to be limited, it is intrinsically interesting to determine whether these cells may represent a potential source of new neurons that may replenish and replace damaged and lost neuronal tissue.

To address this question, it is necessary to understand the intrinsic behavior of NSCs derived from the activated SVZ and SGZ neurogenic niches after TBI, in terms of cell maturation, and differentiation capacity.

In this study, we induced a focal TBI lesion specifically targeting the SVZ or SGZ neurogenic niche in adult rats, harvested NSCs three days post-lesioning, and subsequently expanded them in culture. We found that the isolated NSCs displayed distinct proliferation, differentiation, and spontaneous organization *in vitro*, dependent on the activated niche of origin. Furthermore, these behaviors differed from NSCs derived from the SVZ or SGZ niche of uninjured control animals.

## Introduction

Neural stem cells (NSCs) are multipotent cells capable of self-renewal and differentiation into all neural lineage cells, i.e., neurons, astrocytes, and oligodendrocytes (Bazan et al 2004, Gage 2000, Price & Williams 2001). Within the central nervous system (CNS), NSCs primarily reside within the three canonical neurogenic niches. Two of the canonical neurogenic niches are in the brain, i.e., the subventricular zone (SVZ) of the lateral ventricle and the subgranular zone (SGZ) in the (DG) of the hippocampus. The third canonical neurogenic niche is the central canal (CC) of the spinal cord (Doetsch 2003, Johansson et al 1999, Schiro et al 2022, Taupin 2006). All these niches share a common embryonic mesendodermal developmental origin (Bjornsson et al 2015, Kennea & Mehmet 2002, Redmond et al 2019, Taupin 2006). In the adult mammalian CNS, under physiological conditions, the resident SVZ and SGZ NSCs undergo asymmetric division, resulting in neuronal progeny that migrates and integrates into the neural circuitry in the olfactory bulb and the DG of the hippocampus, respectively (Abbott & Nigussie 2020, Lim & Alvarez-Buylla 2016, Obernier & Alvarez-Buylla 2019, Song et al 2002). The NSCs in the adult mammalian brain neurogenic niches are finely tuned to their specific local microenvironment (Obernier & Alvarez-Buylla 2019), and their activity is regulated by several external factors, such as neuronal and vascular signaling and homeostatic microglial activity (Káradóttir & Kuo 2018, Mizrak et al 2019, Xavier et al 2015, Young et al 2011).

Previous studies have shown that the SGZ niche microenvironment is capable of inducing neuronal differentiation in neuronal progenitors and NSCs transplanted from non-neurogenic environments or neurogenic niches, such as the CC of the spinal cord. These transplanted NSCs differentiate to the point where they can functionally integrate into the local neural circuitry of the hippocampal DG, indistinguishably from SGZ-derived granule cells (Ghosh 2019, Shihabuddin et al 2000). Traumatic disruption to the SGZ microenvironment, such as that caused by TBI, may result in significant, long-lasting, and potentially unpredictable cell behaviors *in vivo*, even significant and prolonged necrotic cell death within the NSC progeny population (Rola et al 2006, Ryu et al 2016, Zhou et al 2012). In response to a traumatic event such as TBI, both neurogenic niches of the brain respond by upregulating NSC proliferative activity, albeit with differentiation capacity towards a glial rather than a neuronal fate (Rola et al 2006). In response to TBI, it is primarily the SVZ NSC progeny that migrates towards the injury site, where they continue to proliferate and differentiate into mature glial cells that subsequently form part of the glial (Salman et al 2004)

The primary focus of this study was to investigate potential lesion-dependent alterations in the survival, proliferative, and differentiation capacity, and self-organization *in vitro* of NSCs isolated from the SVZ and SGZ neurogenic niche, in response to direct, focal TBI-induced disruption of the respective niche microenvironment *in vivo*. To achieve this, we used our optimized protocol (Schiro et al 2022) for the isolation of NSCs from the SVZ and SGZ neurogenic niches to harvest, expand and differentiate NSCs derived from uninjured control and TBI-lesioned animals 3 days post-injury.

As expected, we observed an increase in the proliferative activity of NSCs isolated from both neurogenic niches in response to a TBI lesion. Interestingly, during induced neural differentiation, NSCs harvested from both SVZ uninjured control and TBI conditions generated a large quantity of neural lineage cells that differed greatly in the degree of neuronal versus glial fate and spontaneous structural organization *in vitro*. The uninjured control SVZ-derived NSCs developed small, interconnected clusters, while the TBI-activated SVZ-derived NSCs formed more uniform monolayer neural cultures. We observed the opposite effect in the SGZ-derived NSCs when differentiating, where both uninjured control and TBI lesion-activated NSC cultures formed large colonies of uniform monolayers with a propensity towards a glial fate and very limited capacity for the generation of neuronal cells. We also observed significant cell death during differentiation in the SGZ-derived TBI-activated NSC cultures, reminiscent of the high rate of necrotic cell death previously reported to occur *in vivo*.

## Materials and Methods

### Animals

All animal procedures were approved by the Norwegian Animal Research Authority and were in accordance with the Norwegian Animal Welfare Act §§ 1-28, the Norwegian Regulations of Animal Research §§ 1-26, and the European Convention for the Protection of Vertebrate Animals used for Experimental and Other Scientific Purposes (FOTS ID: 18066 & 24506). The animals were kept on a 12 h light/dark cycle under standard laboratory conditions (19–22°C, 50%–60% humidity), and had *ad libitum* access to food and water. All animals used were 10-week-old female Sprague Dawley rats (Janvier Labs, Le Genest-Saint-Isle, France), housed at the Comparative Medicine core facility, NTNU. A total of 36 animals were used in this study, divided into four groups: Group 1 – SVZ uninjured control: n=9, Group 2 – SVZ TBI-activated: n=9, Group 3 – SGZ uninjured control: n=9, Group 4 – SGZ TBI-activated: n=9.

### Surgical procedure for induction of SVZ and SGZ targeting TBI lesion

Each animal was anesthetized before the surgical procedure and anesthesia was maintained with 2%-3% isoflurane (Baxter, ESDG9623C) in 40% oxygen and 60% nitrogen gas mix, with body temperature regulated by a heating pad beneath the animal. Eye-protective gel (Viscotears, 7939) was applied to the animaĺs eyes immediately after being positioned for surgery. The animal’s head was immobilized in a stereotaxic frame (Harvard apparatus #75-1800) and shaven clean before a local injection of 0.1 ml Marcaine (2,5 mg/ml: Aspen Pharma Trading Limited, Ireland) was administered subcutaneously at the site of surgery as a local analgesic. A longitudinal incision was made along the midline of the skull and the skin was held aside and tissue removed to expose both bregma and lambda. The surgical wound was kept continuously moist using saline (Cat# 190/12606090/0713). Coordinates were defined for SVZ (A/P: -0.6 mm, M/L: +/-4.0 mm, D/V: -4.5 mm) and SGZ (AP: -6.0 mm, ML: +/-5.0 mm, D/V: 4.0 mm) lesion locations (Paxinos & Watson 1997) using a Desktop digital stereotaxic instrument, Dual M (RWD, 68804) connected to the stereotaxic frame for the respective animals before burr holes were drilled through the skull to expose the brain, and a blunted 0.85 mm diameter IV cannula was rapidly lowered manually to the target depth at 1cm/0.5 seconds to simulate a shard of bone/foreign object entering the brain at force. The cannula was held in place for 60 seconds before being carefully removed (0.5 cm/5 seconds) and the surgical wound was closed by sutures.

After the TBI lesions, the animals were allowed to wake up in a heated cage before being transferred to a single-housing cage once consciousness was regained. Over the next 3 days post-surgery, the health of the animals was closely monitored every 6-8 hours. Temgesic 0.45 ml (0.03 mg/ml, Indivor Europe Limited) was administered intraperitoneally as a general analgesic to manage pain for the first 48 hours for all operated animals, and for longer if deemed necessary, based on the Rat Grimace Scale (Sotocina Susana & Sorge Robert 2011).

**Fig. 1:**
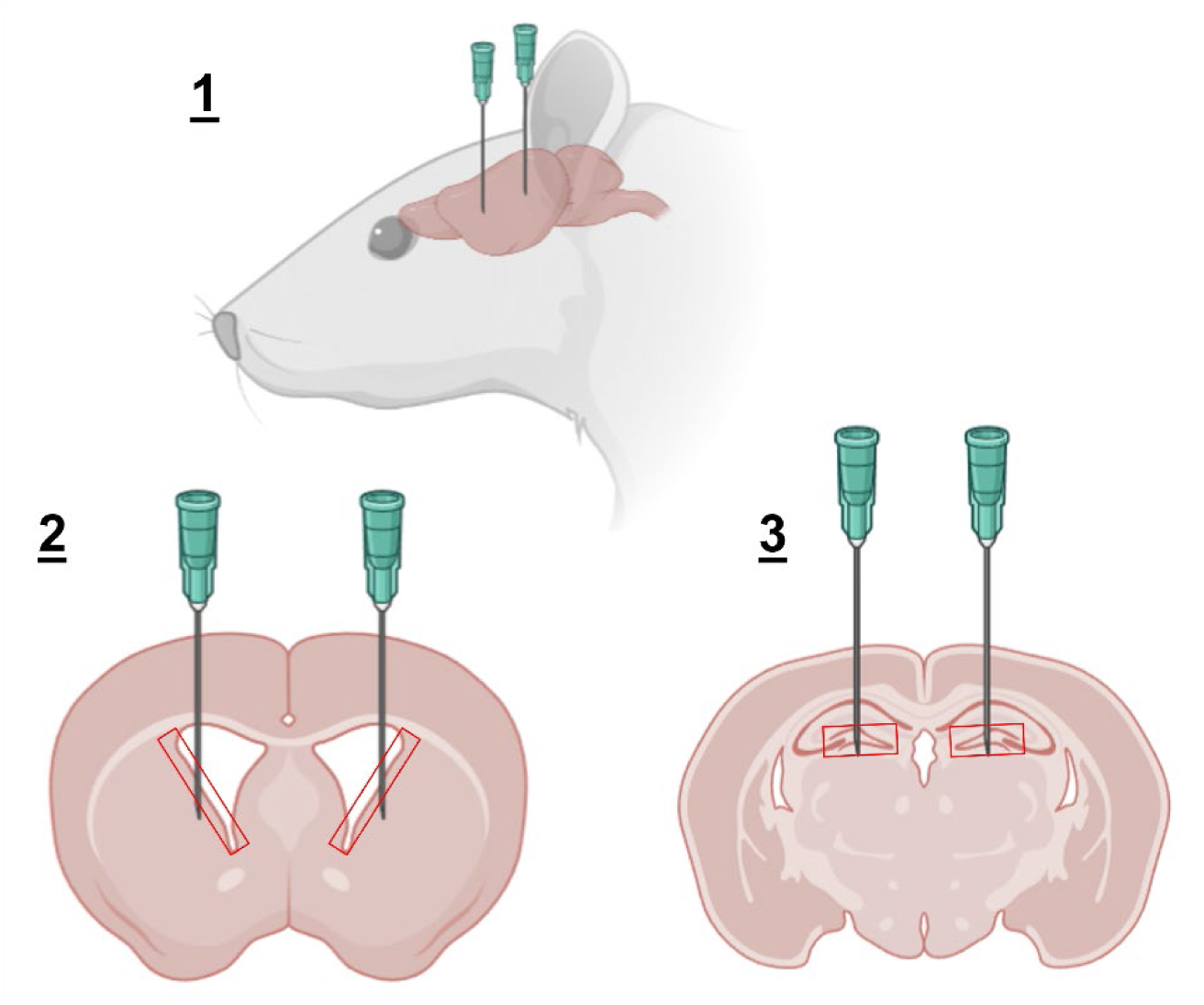
Schematic of SVZ and SGZ lesion and NSC harvest locations. 1: Anterior-posterior locations of SVZ and SGZ lesions. 2: Bilateral TBI lesioning of the subventricular zone neurogenic niche by blunted cannula insertion. Red box: Neurogenic niche harvested. 3: Bilateral TBI lesioning of the subgranular zone neurogenic niche by blunted cannula insertion. Red box: Neurogenic niche harvested.

### Tissue harvest and NSC isolation

The NSCs were isolated from uninjured control and TBI-lesioned animals on the same day. For the TBI-lesioned animals, this was done at 3 days post lesioning, i.e., during the peak of NSC proliferation in response to TBI (Itoh et al 2005, Itoh et al 2007).

The NSCs were expanded for 20 days in expansion media consisting of Neurobasal 1x base medium (Gibco, Cat# 21103049), B27 Without Vitamin A (Gibco, Cat# 12587010), 1x N2 (Gibco, Cat# 17502047), 1x L-Glutamine (Gibco, Cat# 25030081) and Heparin 2.5 µg/mL (Sigma-Aldrich, H3149), supplemented with EGF (Sino biological, Cat# 10605-HNAE), bFGF (Gibco, PHG0026) and CHIR 99021 (Cat# 72052) in sterile non-tissue culture treated 6-well plates (Thermo Fisher Scientific, Cat# 150239). NSCs were passaged using either accutase (Stem cell technologies, Cat# 07920) or trypsin (Thermo Fisher, Cat# 25200056) and counted on days 10 and 20, with all SGZ-derived NSCs transferred to Sterile tissue culture treated 6-well plate (VWR, Cat# 10861-696) at 10 DIV following P1 passage. The NSCs were seeded on Poly-L-Ornithine (Sigma-Aldrich, P4957) and Natural Mouse Laminin (Gibco, Cat# 23017-015) coated coverslips after P2 passage, and expanded for 3 days (23 DIV) before either fixation with a 3% glyoxal solution (Richter et al 2018) or differentiation. A subset of NSC cultures was maintained in an EdU-supplemented expansion media for 24 hours before fixation (22 DIV) for labeling of proliferating cells.

Detailed protocols for NSC isolation and expansion, EdU labeling, tissue processing, and immunocytochemistry can be found in (Schiro et al 2022). Antibodies used are listed in S1.

### Tissue processing and immunohistochemistry

For labeling of proliferating cells in the CC neurogenic niche after injury, each animal was injected intraperitoneally with 37 mg/kg BW of 5-ethynyl-2′-deoxyuridine (EdU, ThermoFisher, A10044) at 48 hours after surgery, and kept alive for 24 hours before tissue fixation by intracardial perfusion. The animals were anesthetized with 4% isoflurane before an overdose of Pentobarbital (5 ml/kg Bodyweight, Ås Produksjonslab AS, 005608) administered by intraperitoneal injection. Each animal was then perfused by an initial 50 ml of heparin-supplemented saline (1:1000, LEO Pharma, 09837) solution at room temperature (23°C) at a rate of 20 ml/min, followed by 200 ml of chilled (4°C) 4% fresh paraformaldehyde (PFA, Sigma, P6148) at 20 ml/min. Once the perfusion was complete, the spinal cord was extracted and post-fixated overnight in 4% PFA at 4°C, before being transferred to a cryoprotective sinking solution (30% Sucrose (Sigma, 84097) in MQ water) and stored for 1 week at 4°C before sectioning. The spinal cords were sectioned coronally at 40 µm in 6 equally spaced series on a freezing microtome for immunolabelling and analysis.

One randomly selected series of tissue sections from each spinal cord were dehydrated in ethanol, cleared in xylene (VWR International, Radnor, PA, USA, 1330-20-7), and rehydrated before staining with Cresyl violet (Nissl; Sigma-Aldrich, St. Louis, MO, USA, C5042-10G) for 3 minutes on a shaker while protected from light. The sections were then alternatively dipped in ethanol–acetic acid (5 ml acetic acid in 1L 70 % ethanol) and rinsed with cold water until the desired differentiation was obtained, then dehydrated, cleared in xylene, and coverslipped with Entellan containing Xylene (VWR International, Radnor, PA, USA, 100503-868). For EdU labeling, one series of tissue sections from each spinal cord was incubated in reaction buffer (Click-IT), CuSO4, Alexa Fluor Azide (488), and reaction buffer additive (Invitrogen, C10337) for 30 minutes. The incubation was followed by a wash with 4′, 6-diamidino-2-fenylindol (DAPI; 1:10 000; Sigma-Aldrich, Saint-Louis, MO, USA, 28718-90-3) and phosphate buffer (PB) for 10 minutes, followed by a wash in PB for 10 minutes.

For immunohistochemistry, heat-induced antigen retrieval (HIAR) was carried out on two series of tissues (stained with Nestin/Ki67 and Iba1/GFAP) at 60 °C for 2 hours in PB. Sections were subsequently washed 3 times for 10 minutes with PB containing 0.2 % Triton X-100 (Merck, Darmstadt, Germany, 108603; PBT). Next, sections were blocked using 5 % normal goat serum (Abcam, Cambridge, UK, ab7481) in PBT (PBT+) for 1 hour before incubation with the primary antibody in PBT+ for 4 hours at 4 °C (see table S1 for primary antibodies used). Sections were subsequently washed 3 times for 10 minutes in PBT followed by incubation with Alexa Fluor secondary antibodies (Invitrogen, California, USA) at a 1:400 concentration (Alexa Fluor 488, 568, and 647, Table S1) for 2 hours at room temperature, protected from light. Then sections were incubated for 10 minutes with DAPI (1:10 000) in PB, followed by 3 washes for 10 minutes in PB.

### Differentiation of SVZ and SGZ-derived NSCs

All NSC cultures were differentiated for a total of 30 days, with 80% of media changed every 3 days. The cells were imaged after every passage and media change to track development. Differentiation base media: Neurobasal 1x base medium (Gibco, Cat# 21103049), 1x B27 Plus (Gibco, A3582801), 1x N2 (Gibco, Cat# 17502047), 1x L-Glutamine (Gibco, Cat# 25030081) and Heparin 2.5 µg/mL (Sigma-Aldrich, H3149). Differentiation supplement SVZ and SGZ: BDNF (PeproTech, Cat# 450-02). The differentiated cultures were fixed after 30 days in differentiation media, fixed and immunolabelled according to the protocol published in (Schiro et al 2022).

### Imaging and quantification of cell proliferation and survival

Live cells were imaged using a Zeiss Axiovert 25 inverted phase contrast microscope with a Zeiss Axiocam 105 color camera and transmitted white light as the light source.

Cell proliferation and survival: Total cell counts, and percentage of live cells were counted automatically following dissociation by staining the cells with Trypan blue and counting using a Countess™ II Automated Cell Counter (Invitrogen, AMQAX1000) and averaged for each animal after each passage.

Tissue sections were imaged using a Mirax-midi slide scanner and an Axiovert A1 fluorescent microscope (Zeiss, 503 mono camera) using either reflected fluorescence (fluorescence imaging) or transmitted white light (Cresyl Violet; Nissl) as the light source.

Immunolabelled cell cultures were imaged using an EVOS M5000 (Thermo Fisher) and counted using FIJI’s (ImageJ) automated cell counting function based on ICC images with a 1.0 mm^2^ field of view per image. Structural network nodes and connecting filaments were counted manually from random 1 mm^2^ sample images from each differentiated culture.

Statistical data was tested for normal distribution (Shapiro-Wilk) and equal variance (F-test (Snedecor & Cochran 1989)) before mean comparison using two-tailed unpaired t-test for normally distributed and small sample size data (n < 6 (De Winter 2019)) or Mann Whitney U for non-parametric data in STATA/MP 18.

## Results

### Histological verification of TBI lesion and lesion-induced NSC activation *in vivo*

The SVZ and SGZ neurogenic niches were histologically identified by Nissl staining (S4A1-2). Inflammatory response to TBI lesion was verified by the presence of Iba-1 positive microglia within brain sections from SVZ and SGZ TBI lesioned animals, with none observed in the uninjured control derived brain sections (S4B&C1-2). Nestin-positive cells were primarily located within the SVZ and SGZ neurogenic niches in the uninjured control brain sections (S4D1-2). However, Nestin-positive cells were observed within the tissue surrounding the neurogenic niches and lesion sites in brain sections derived from both SVZ and SGZ TBI lesioned animals (S4E1-2).

### Expansion of NSCs as adherent monolayer cultures

A total of 500,000 live cells per neurogenic niche were seeded for expansion for each experimental condition and cultured as a cell suspension in sterile non-tissue culture-treated 6-well plates to encourage the NSCs to form free-floating neurospheres (Fig. 2A-D1, red arrows). The free-floating neurospheres were allowed to spontaneously settle on the culture vessel surface, where they formed monolayer colonies at 4-7 DIV (Fig. 2A-D2). This pattern of NSC development was consistent across all cultures and conditions after P1 dissociation and reseeding as single-cell suspensions at 10 DIV (Fig. 2A-D3).

**Fig. 2:**
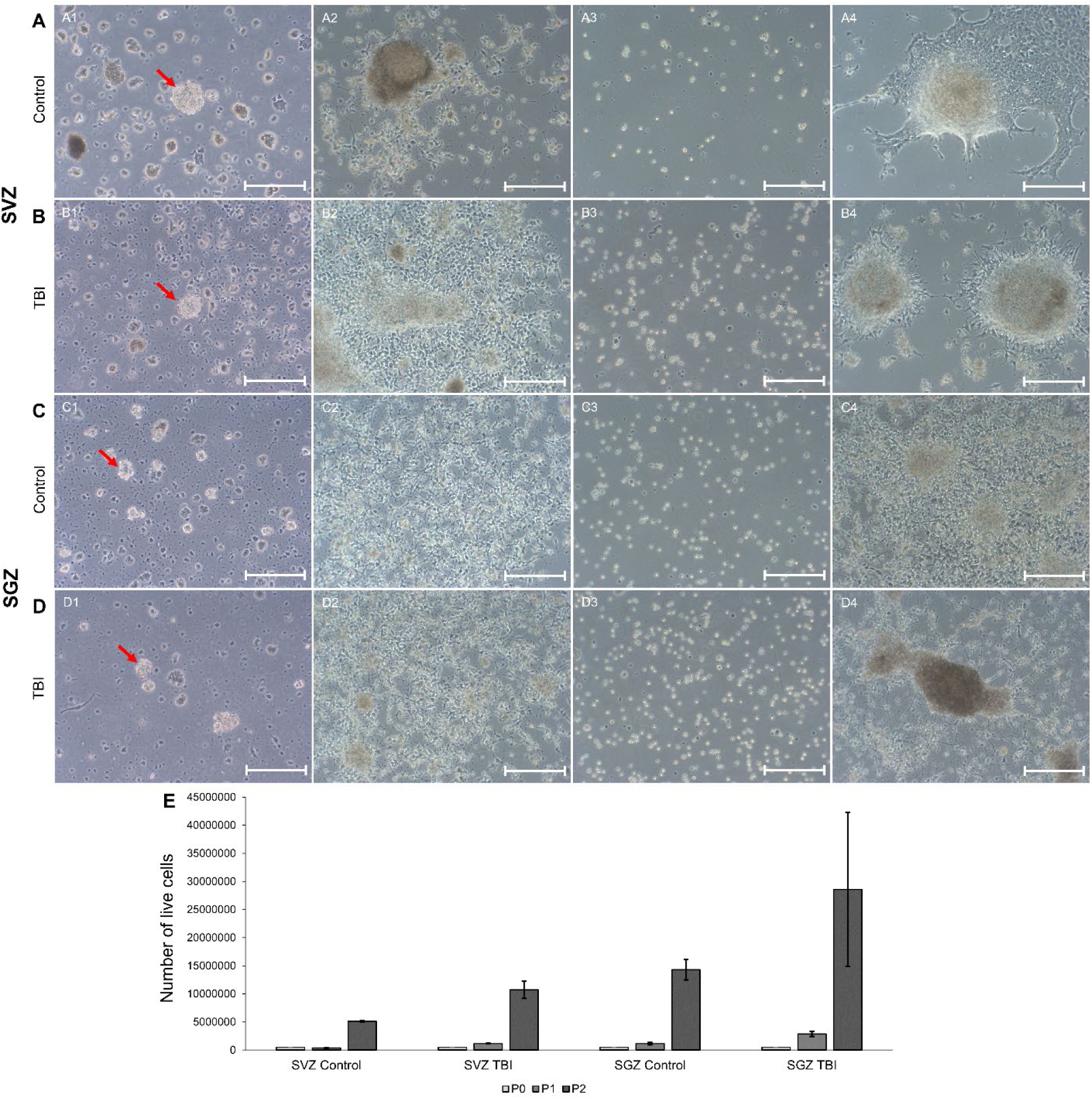
SVZ and SGZ-derived uninjured control and TBI-activated NSC expansion over 20 days *in vitro*. Brightfield images of SVZ and SGZ uninjured control and TBI-activated NSC culture development during expansion. A1: Free floating neurosphere (red arrow) of SVZ uninjured control NSC cultures at 4 DIV. A2: Pre-P1 passage: monolayer colony of SVZ uninjured control NSC cultures at 10 DIV. A3: Post P1 passage: monolayer colonies of SVZ uninjured control NSC cultures at 10 DIV. A4: Pre-P2 passage: monolayer colony of SVZ uninjured control NSC cultures at 20 DIV. B1: Free floating neurosphere (red arrow) of SVZ TBI-activated NSC cultures at 4 DIV. B2: Pre-P1 passage monolayer colony of SVZ TBI-activated NSC cultures at 10 DIV. B3: Post P1 passage: monolayer colony of SVZ TBI-activated NSC cultures at 10 DIV. B4: Pre-P2 passage: monolayer colony of SVZ TBI-activated NSC cultures at 20 DIV. C1: Free floating neurosphere (red arrow) of SGZ uninjured control NSC cultures at 4 DIV. C2: Pre-P1 passage: monolayer colony of SGZ uninjured control NSC cultures at 10 DIV. C3: Post P1 passage: monolayer colony of SGZ uninjured control NSC cultures at 10 DIV. C4: Pre-P2 passage: monolayer colony of SGZ uninjured control NSC cultures at 20 DIV. D1: Free floating neurosphere (red arrow) of SGZ TBI-activated NSC cultures at 4 DIV. D2: Pre-P1 passage: monolayer colony of SGZ TBI-activated NSC cultures at 10 DIV. D3: Post P1 passage: monolayer colony of SGZ TBI-activated NSC cultures at 10 DIV. D4: Pre-P2 passage: monolayer colony of SGZ TBI-activated NSC cultures at 20 DIV. All scale bars: 200 µm. E: NSCs harvested from all conditions displayed a significant increase in proliferative activity between 10- and 20-days post-harvest, regardless of injury status (uninjured control or TBI-activated), with only the SVZ TBI-activated NSC cultures displaying significantly increased proliferative activity compared to the SVZ uninjured control NSC cultures at P2. See Table S2 for expanded statistics.

A slight reduction in the number of live cells in the SVZ uninjured control NSC cultures was observed at P1 (Fig. 2A2), followed by a significant (t_2_ = 49.320, p = 0.0004, unpaired two-tailed t-test) 13.2-fold increase in live cells by P2 (Fig. 2A4&E, Table S2).

A 2.36-fold increase in the total number of live cells (Fig. 2B2) from P0 to P1 was observed across the SVZ TBI-activated NSC cultures, a significantly greater increase (t_2_ = 9.905, p = 0.010, unpaired two-tailed t-test) compared to the SVZ uninjured control NSC cultures. The number of live cells in the SVZ TBI-activated NSC cultures further increased 9.08-fold from P1 to P2 (Fig. 2B4), resulting in higher, but not statistically significant (t_2_ = 5,171, p = 0.119, unpaired two-tailed t-test) number of live cells at P2 compared to the SVZ uninjured control NSC cultures (Fig. 2E, Table S2).

The number of live cells derived from the SGZ uninjured control NSC cultures increased 2.33-fold at P1 compared to P0 (Fig. 2C2), a significantly greater increase than what was observed in the SVZ uninjured control NSC cultures (t_4_ = 4.386, p = 0.0118, unpaired two-tailed t-test). This increase in total live cells was however comparable to that of the SVZ TBI-activated NSC cultures (t_4_ = 0.068, p = 0.948, unpaired two-tailed t-test). The number of live cells in the SGZ uninjured control NSC cultures further increased 12.24-fold from P1 to P2 (Fig. 2C4). A significant increase compared to P2 SVZ uninjured control NSC cultures (t_4_ = 9.939, p = 0.002, unpaired two-tailed t-test), and comparable to the increase of the SVZ TBI-activated NSC cultures at P2 (t_4_ = 2.339, p = 0.079, unpaired two-tailed t-test, Fig. 2E, Table S2).

In the SGZ TBI-activated NSC cultures, we observed a significantly (t_6_ = 6.900, p = 0.0005) larger initial 5.72-fold increase in the number of live cells between P0 to P1 (Fig. 2D2) compared to the SGZ uninjured control, the SVZ uninjured control (t_4_ = 7.598, p = 0.0016) and SVZ TBI-activated NSC cultures (t_4_ = 5.157, p = 0.0067). We observed a 9.98-fold increase in the number of live cells from P1 to P2 in the SGZ TBI-activated NSC cultures (Fig. 2D4), resulting in a comparable (t_6_ = 2.064, p = 0.127) number of live cells to the SGZ uninjured control, and SVZ TBI-activated NSC cultures (t_4_ = 1.733, p = 0.158). In comparison, the increase in live cells in the SGZ TBI-activated NSC cultures was significantly (t_4_ = 3.423, p = 0.0417) greater than the number of live cells from the SVZ uninjured control NSC cultures (Fig. 2E, Table S2).

### Distinct NSC proliferative and differentiation patterns were revealed in response to TBI

After P2 passage (20 DIV) the NSCs from the uninjured control animals and NSCs derived from SVZ and SGZ TBI lesioned animals were seeded on PLO/Laminin coated 13 mm^2^ glass coverslips (13 mm^2^) at 500 000 cells per condition per coverslip and expanded for another 3 days before fixation (Fig. 3A-D1).

**Fig. 3:**
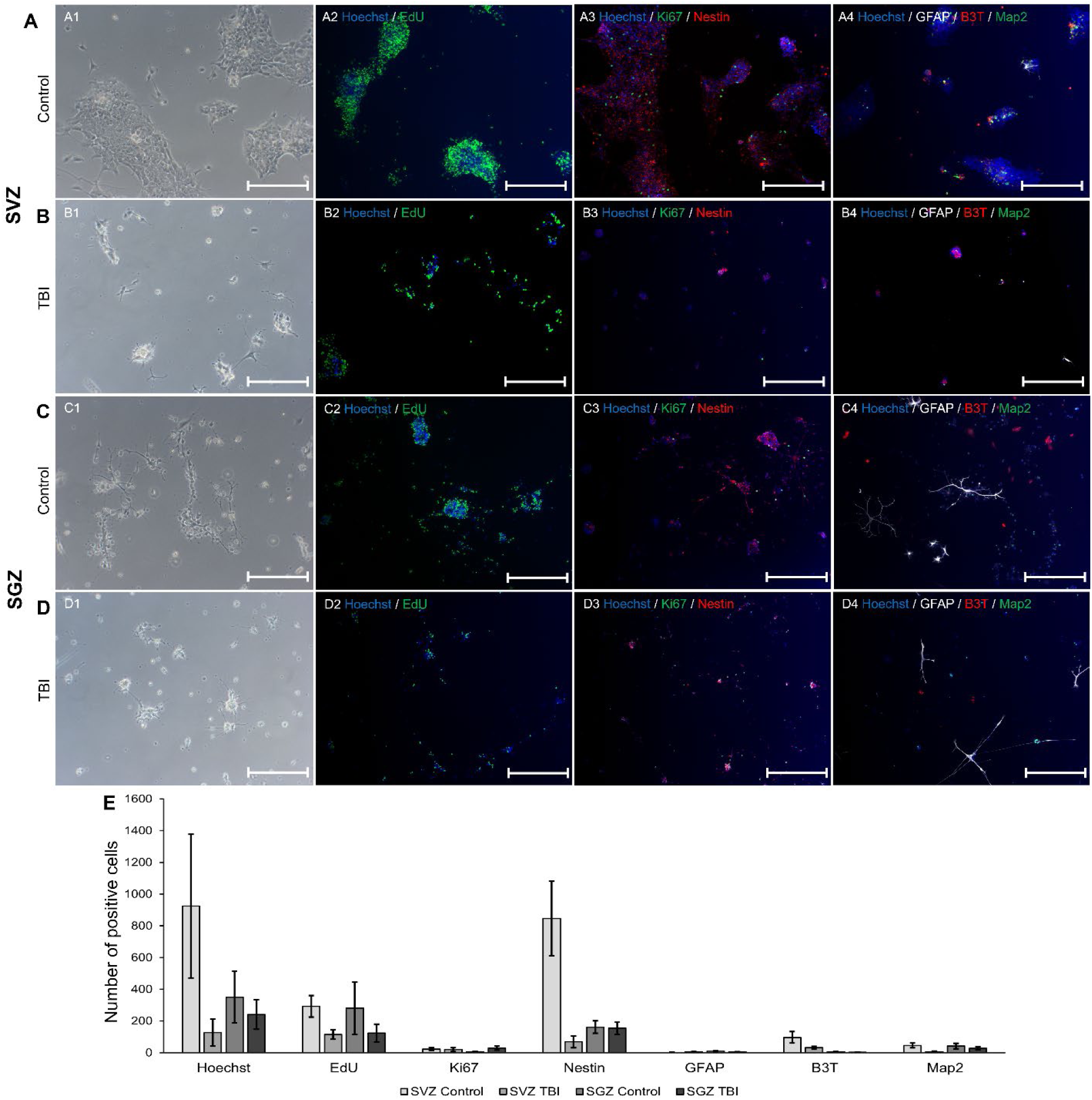
Expression of proliferation and NSC markers after expansion. A1: SVZ uninjured control NSC cultures after 3 days on PLO/laminin-coated coverslips before fixation (23 DIV). A2: EdU labeling of proliferating SVZ uninjured control NSC cultures after 24-hour EdU incubation before fixation (22-23 DIV). A3: Actively proliferating (Ki67) SVZ uninjured control NSCs (Nestin). A4: SVZ uninjured control NSC cultures neural (GFAP, Beta-III-Tubulin (B3T) and Map2) spontaneous differentiation at 23 DIV. B1: SVZ TBI-activated NSC cultures after 3 days on PLO/laminin-coated coverslips before fixation (23 DIV). B2: EdU labeling of proliferating SVZ TBI-activated NSC cultures after 24-hour EdU incubation before fixation (22-23 DIV). B3: Actively proliferating (Ki67) SVZ TBI-activated NSCs (Nestin). B4: SVZ TBI-activated NSC cultures neural (GFAP, Beta-III-Tubulin (B3T) and Map2) spontaneous differentiation at 23 DIV. C1: SGZ uninjured control NSC cultures after 3 days on PLO/laminin-coated coverslips before fixation (23 DIV). C2: EdU labeling of proliferating SGZ uninjured control NSC cultures after 24-hour EdU incubation before fixation (22-23 DIV). C3: Actively proliferating (Ki67) SGZ uninjured control NSCs (Nestin). C4: SGZ uninjured control NSC cultures neural (GFAP, Beta-III-Tubulin (B3T) and Map2) spontaneous differentiation at 23 DIV. D1: SGZ TBI-activated NSC cultures after 3 days on PLO/laminin-coated coverslips before fixation (23 DIV). D2: EdU labeling of proliferating SGZ TBI-activated NSC cultures after 24-hour EdU incubation before fixation (22-23 DIV). D3: Actively proliferating (Ki67) SGZ TBI-activated NSCs (Nestin). D4: SGZ TBI-activated NSC cultures neural (GFAP, Beta-III-Tubulin (B3T) and Map2) spontaneous differentiation at 23 DIV. All scale bars: 200 µm. E: Following expansion both the SVZ uninjured control and SGZ TBI-activated NSCs cultures indicated stable cultures with a slower turnover rate, as indicated by Ki67 and EdU labeling. In contrast, the SVZ TBI-activated and SGZ uninjured control NSC cultures both displayed a high turnover with 90.2% and 80.1% EdU labeled Hoechst positive nuclei after 24h incubation respectively. Spontaneous neural differentiation at 23 DIV was low across all NSC cultures from both neurogenic niches and experimental conditions (uninjured control and TBI-activated). See Table S3 for expanded statistics.

ICC labeling of the SVZ uninjured control NSC cultures revealed an average of 923.8 cells per 1 mm^2^ as indicated by Hoechst positive nuclei. 31.6% of the total cell population had EdU positive nuclei after a 24-hour incubation period (Fig. 3A2), with 2.5% of the total cell population undergoing S-phase cell division, as indicated by the number of Ki67 positive cell nuclei at the time of fixation (Fig. 3A3). We found that 91.6% of the total cell population was positive for the NSC marker Nestin (Fig. 3A3). Only a small subset of cells underwent spontaneous neural differentiation, with 0.14% of the total cell population being positive for GFAP, 10.4% positive for Beta-III-Tubulin, and 4.9% of the total cell population positive for Map2 (Fig. 3A4&E, Table S3).

In the TBI-activated SVZ NSC cultures, we observed an average of 127.1 Hoechst positive cell nuclei per 1 mm^2^, significantly lower than the uninjured control cultures (z_58_ = 6.653, p < 0.0001, Mann Whitney U). However, a significantly higher proportion of the Hoechst positive cell nuclei was EdU labeled after 24-hour incubation (90.2%), indicating a significantly higher cell turnover rate in the SVZ TBI-activated NSC cultures compared to the SVZ uninjured control NSC cultures (z_16_ = 3.554, p = 0.0004, Mann Whitney U, Fig. 3B2). 15.98% of total cell population was found to undergo S-phase cell division at the time of fixation (Fig. 3B3), indicating a similar, relative turnover rate compared to the SVZ uninjured control NSC cultures (z_18_ = 6.653, p < 0.0001, Mann Whitney U). A significantly lower proportion of the total SVZ TBI-activated NSC cell population (53.7%, z_17_ = 3.676, p = 0.0002, Mann Whitney U, Fig. 3B3) was found to be Nestin positive compared to the SVZ uninjured control NSC cultures. A significantly larger proportion of TBI-activated SVZ NSC cultures underwent spontaneous neural differentiation compared to the SVZ uninjured control NSC cultures with 4.4% of the total cell population positive for GFAP (z_18_ = 3.509, p = 0.0004, Mann Whitney U), 25% Beta-III-Tubulin (z_18_ = 3.781, p = 0.0002, Mann Whitney U), however, the proportion of Map2 (4.7%) positive cells was comparable between the two conditions (z_18_ = 0.606, p = 0.544, Mann Whitney U), Fig. 3B4&E, Table S3).

In the SGZ uninjured control NSC cultures, we found an average of 349.9 Hoechst-positive nuclei per 1 mm^2^, significantly less than the SVZ uninjured control (z_58_ = 6.150, p < 0.0001, Mann Whitney U) and SVZ TBI-activated (z_58_ = 5.752, p < 0.0001, Mann Whitney U) NSC cultures. An 80.1% co-labeling of Hoechst positive nuclei with EdU after a 24-hour incubation period indicated a high relative turnover rate, significantly higher than that of the SVZ uninjured control NSC cultures (z_18_ = 3.326, p = 0.0009, Mann Whitney U), but comparable to the SVZ TBI-activated NSC cultures (z_16_ = 0.978, p = 0.328, Mann Whitney U, Fig. 3B2). However, we found that 1.6% of the total cell population underwent S-phase cell division at the time of fixation, significantly lower than both SVZ uninjured control (z_18_ = 2.425, p = 0.015, Mann Whitney U) and SVZ TBI-activated NSC cultures (z_18_ = 3.790, p = 0.0002, Mann Whitney U). 46% of the total cell population was Nestin positive, significantly lower than the SVZ uninjured control (z_17_ = 3.511, p = 0.0004, Mann Whitney U), but comparable to the SVZ TBI-activated NSC cultures (z_18_ = 0.227, p = 0.820, Mann Whitney U, Fig. 3C3). A small proportion of the SGZ uninjured control NSC cultures underwent spontaneous neural differentiation with 2.6% positive for GFAP, a significantly higher proportion than the SVZ uninjured control (z_18_ = 3.801, p = 0.0001, Mann Whitney U), but significantly lower than the SVZ TBI-activated NSC cultures (z_18_ = 2.571, p = 0.0099, Mann Whitney U), 1.6% Beta-III-Tubulin (Significantly lower than both the SVZ uninjured control (z_18_ = 3.788, p = 0.0002, Mann Whitney U) and SVZ TBI-activated conditions (z_18_ = 3.790, p = 0.0002, Mann Whitney U)) and 12.0% Map2 (significantly higher than the SVZ uninjured control (z_18_ = 3.026, p = 0.0025, Mann Whitney U) and the SVZ TBI-activated NSC cultures (z_18_ = 3.180, p = 0.0055, Mann Whitney U, Fig. 3C4&E, Table S3).

We observed an average of 239.8 cells per 1 mm^2^ in the SGZ TBI-activated NSC cultures, significantly lower than both the SGZ and SVZ uninjured control (z_58_ = 3.526, p = 0.0004 and z_58_ = 6.623, p < 0.0001 respectively, Mann Whitney U), but significantly higher than the SVZ TBI-activated NSC cultures (z_58_ = 4.761, p < 0.0001). Of the total cell population 51.7% of the cells were labeled with EdU after a 24-hour incubation period (comparable to the SGZ uninjured control (z_18_ = 1.739, p = 0.0820, Mann Whitney U), but significantly higher than the SVZ uninjured control (z_18_ = 2.344, p = 0.019, Mann Whitney U) and lower than the SVZ TBI-activated NSC cultures (z_16_ = 2.935, p = 0.003, Mann Whitney U, Fig. 3B), alongside 12.0% active S-phase dividing cells (significantly higher than both the SGZ and SVZ uninjured controls (z_18_ = 3.788, p = 0.0002 and z_18_ = 3.781, p = 0.0002, respectively, Mann Whitney U), but comparable to the SVZ TBI-activated NSC cultures (z_18_ = 0.983, p = 0.325, Mann Whitney U) indicate a slower turnover rate than both the SVZ and SGZ uninjured control NSC cultures (Fig. 3D3). 64% of the total cell population was Nestin positive, significantly higher than the SGZ uninjured control (z_18_ = 2.419, p = 0.015, Mann Whitney U) and the SVZ TBI-activated NSC cultures (z_18_ = 1.966, p = 0.043, Mann Whitney U) but significantly lower than the SVZ uninjured control (z_18_ = 2.449, p = 0.014, Mann Whitney U, Fig. 3D3). 1.6% of total cell population spontaneously differentiated to GFAP positive cells, significantly lower than the SGZ uninjured control and SVZ TBI-activated NSC cultures (z_18_ = 2.427, p = 0.015 and z_18_ = 3.493, p = 0.0005 respectively, Mann Whitney U), but significantly higher than the SVZ uninjured control NSC cultures (z_18_ = 3.792, p = 0.0002, Mann Whitney U). Spontaneous differentiation into Beta-III-Tubulin positive cells occurred in 0.95% of the total cell population, a proportion significantly lower than the SGZ (z_18_ = 2.596, p = 0.009, Mann Whitney U) and SVZ uninjured controls (z_18_ = 3.808, p = 0.0001, Mann Whitney U) and SVZ TBI-activated NSC cultures (z_18_ = 3.810, p = 0.0001, Mann Whitney U). Finally, 11.7% of the total SGZ TBI-activated NSC culture cell population spontaneously differentiated into Map2 positive cells, a level comparable to that of the SGZ uninjured control (z_18_ = 0.151, p = 0.879, Mann Whitney U), but significantly higher than both SVZ uninjured control (z_18_ = 3.254, p = 0.001, Mann Whitney U) and TBI-activated conditions (z_18_ = 3.105, p = 0.0019, Mann Whitney U, Fig. 3D4&E, Table S3).

### TBI impacts neural lineage differentiation and spontaneous neural network self-organization *in vitro*

After 3 days of further expansion (23 DIV) on PLO/Laminin coated coverslips NSCs from all culture conditions settled as monolayer colonies, with the occasional multilayered clusters spread sporadically throughout the culture (Fig. 4A-D1), before switching media from expansion to differentiation.

**Fig. 4:**
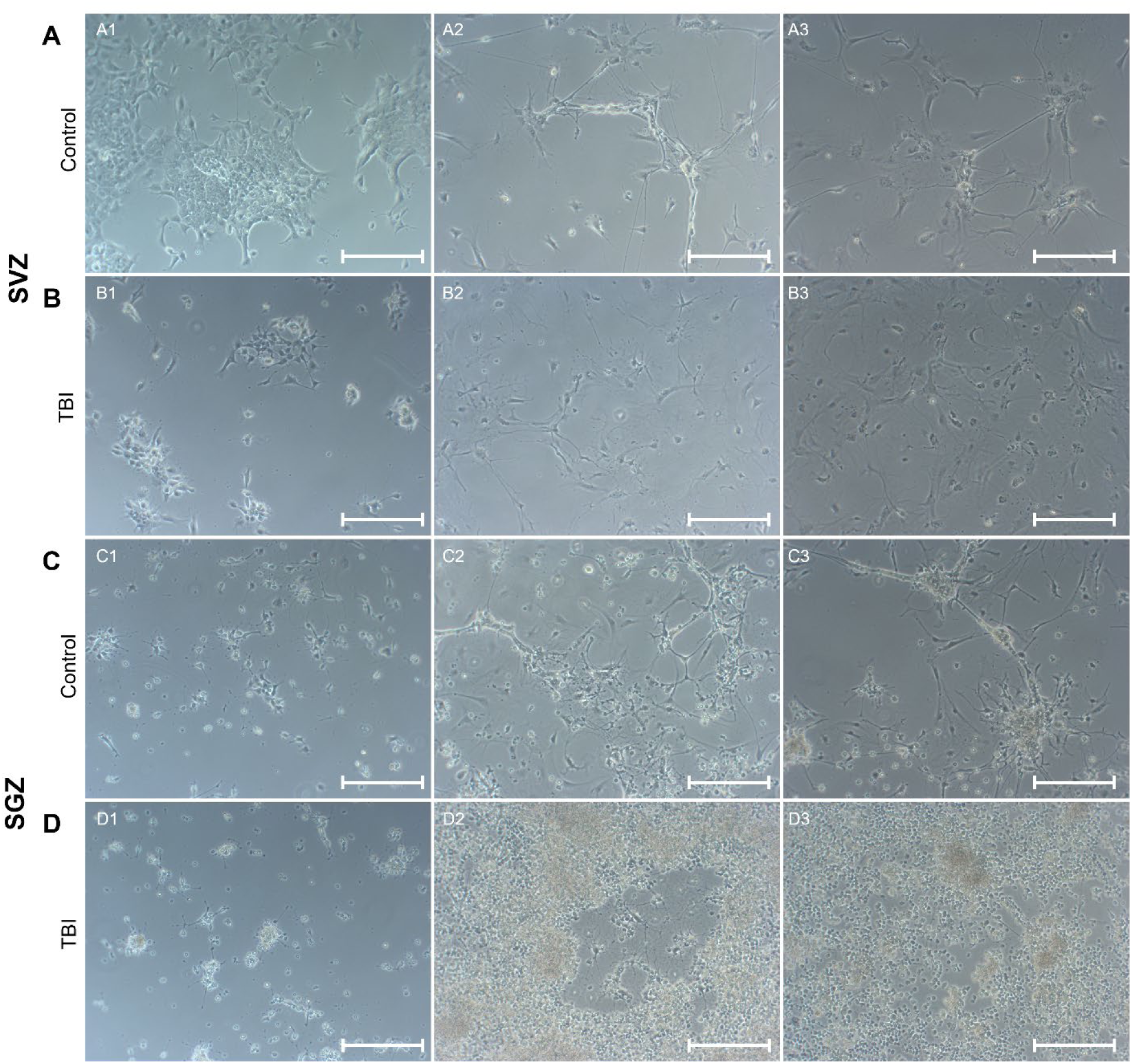
Differentiation and spontaneous self-organization of uninjured control and TBI-activated SVZ and SGZ NSCs over 30 days. Brightfield images of A1: SVZ uninjured control NSC cultures at the start of neural differentiation (23 DIV). A2: SVZ uninjured control NSC cultures at 15 days of neural differentiation (38 DIV). A3: SVZ uninjured control NSC cultures at 30 days of neural differentiation, before fixation (53 DIV). B1: SVZ TBI-activated NSC cultures at the start of neural differentiation (23 DIV). B2: SVZ TBI-activated NSC cultures at 15 days of neural differentiation (38 DIV). B3: SVZ TBI-activated NSC cultures at 30 days of neural differentiation, before fixation (53 DIV). C1: SGZ uninjured control NSC cultures at the start of neural differentiation (23 DIV). C2: SGZ uninjured control NSC cultures at 15 days of neural differentiation (38 DIV). C3: SGZ uninjured control NSC cultures at 30 days of neural differentiation, before fixation (53 DIV). D1: SGZ TBI-activated NSC cultures at the start of neural differentiation (23 DIV). D2: SGZ TBI-activated NSCs at 15 days of neural differentiation (38 DIV). D3: SGZ TBI-activated NSC cultures at 30 days of neural differentiation, before fixation (53 DIV). All scale bars: 200 µm.

After 15 days of differentiation (dDiff) the SVZ uninjured control NSC cultures spontaneously organized into groups of smaller colonies with a subsection of the cells within each colony adopting neuron-like morphology extending processes between colonies (Fig. 4A2). This structural organization of the differentiating NSC cultures remained largely unchanged up to 30 dDiff, with other subsets of cells adopting glia-like morphology. At 30 dDiff the differentiated SVZ uninjured control NSC cultures settled into stable interconnected colonies consisting primarily of Map2, Beta-III-Tubulin, and neurofilament heavy positive cells situated within larger clusters of vimentin-positive but GFAP negative cells (30 dDiff, Fig. 4A3, 5A1-4).

SVZ TBI-activated NSC cultures settled and expanded similarly to the SVZ uninjured control NSC cultures, albeit forming smaller clusters before the start of differentiation (23 DIV, 0 dDiff, Fig. 4 B1). The SVZ TBI-activated NSC cultures spread out into large monolayers by 15 dDiff, with a subset of the cells adopting glia-like morphology with single morphologically neuron-like cells scattered sporadically throughout the cultures (Fig. 4B2). At 30 dDiff large glia-like monolayer colonies of vimentin-positive cells covered approximately 65% of the culture vessel surface with Beta-III-Tubulin and Map2 positive neuronal and GFAP positive glial cells loosely clustered together throughout the cultures. Large solitary neurofilament heavy positive cells with distinct neuronal morphologically extended processes throughout the vimentin-positive cell clusters (Fig. 4B3, 5B1-4).

The SGZ uninjured control NSC cultures settled primarily into small clusters, with a subset of cells settling into unstructured groupings along the external processes of large GFAP positive glial cells (23 DIV, 0 dDiff, Fig. 4C1, 4C4, S5 (Schiro et al 2022)). After 15 dDiff the NSCs had spontaneously organized into large, semi-structured, and interconnected colonies consisting of cells displaying a neuronal-like morphology layered onto a monolayer of large, morphologically glial-like cells at approximately 75% confluence (Fig. 4C2). At 30 dDiff the semi-structured colonies further refined into interconnected colonies of GFAP and vimentin-positive glial cells, with singular Beta-III-Tubulin and Map2 positive neuronal cells embedded within and extending out of a small subset of the glial cell clusters (Fig. 4C3, 5C1-4).

**Fig. 5:**
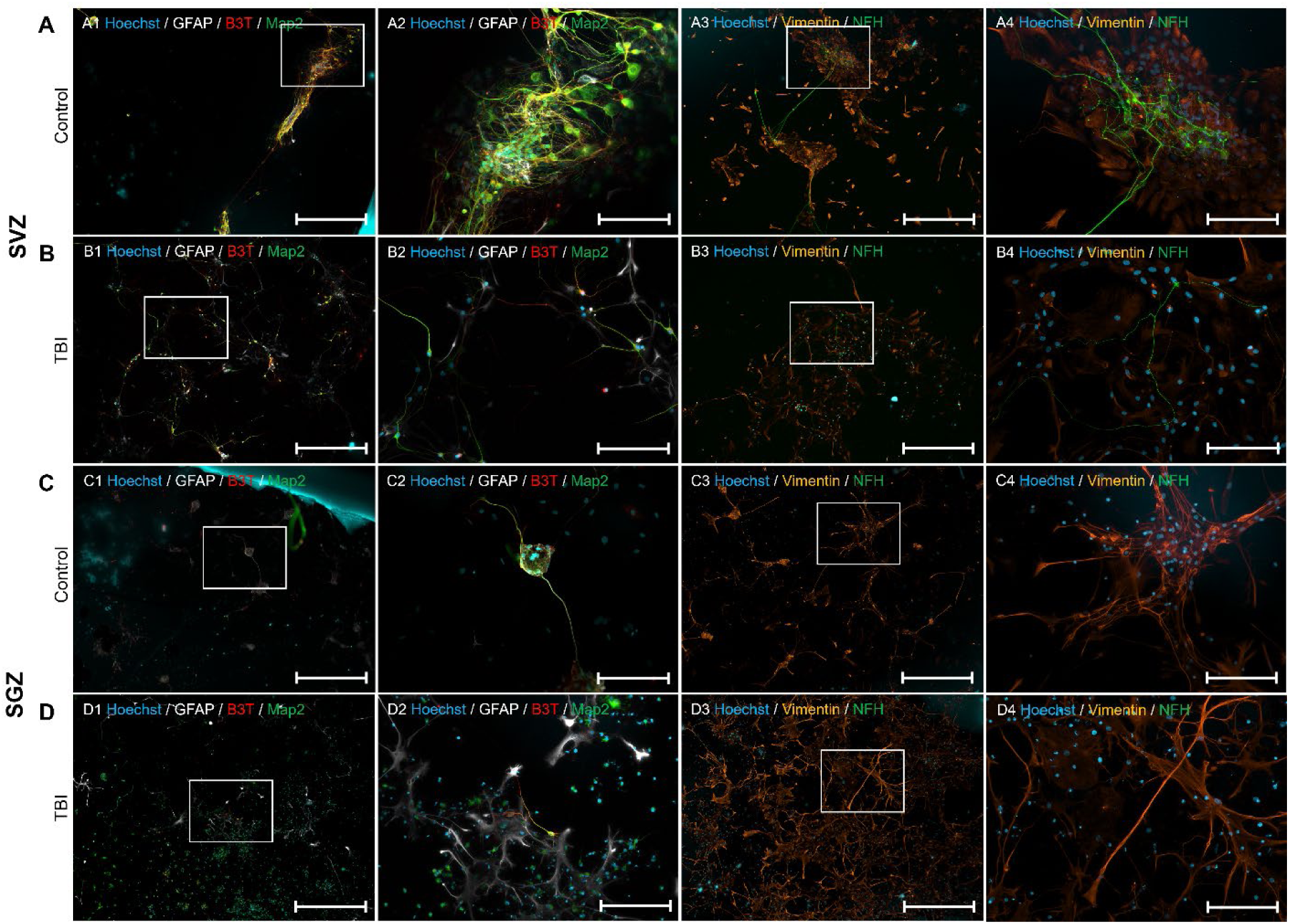
Immunocytochemistry of differentiated uninjured control and TBI-activated SGZ and SVZ NSCs. A1: GFAP, Beta-III-Tubulin, and Map2 immunoreactivity of SVZ uninjured control NSC cultures after 30 days of differentiation. A2: GFAP, Beta-III-Tubulin, and Map2 immunoreactivity of SVZ uninjured control NSC cultures after 30 days of differentiation (A1 white box). A3: Vimentin and neurofilament heavy immunoreactivity of SVZ uninjured control NSC cultures after 30 days of differentiation. A4: Vimentin and neurofilament heavy immunoreactivity of SVZ uninjured control NSC cultures after 30 days of differentiation (A3 white box). B1: GFAP, Beta-III-Tubulin, and Map2 immunoreactivity of SVZ TBI-activated NSC cultures after 30 days of differentiation. B2: GFAP, Beta-III-Tubulin, and Map2 immunoreactivity of SVZ TBI-activated NSC cultures after 30 days of differentiation (B1 white box). B3: Vimentin and neurofilament heavy immunoreactivity of SVZ TBI-activated NSC cultures after 30 days of differentiation. B4: Vimentin and neurofilament heavy immunoreactivity of SVZ TBI-activated NSC cultures after 30 days of differentiation (B3 white box). C1: GFAP, Beta-III-Tubulin, and Map2 immunoreactivity of SGZ uninjured control NSC cultures after 30 days of differentiation. C2: GFAP, Beta-III-Tubulin, and Map2 immunoreactivity of SGZ uninjured control NSC cultures after 30 days of differentiation (C1 white box). C3: Vimentin and neurofilament heavy immunoreactivity of SGZ uninjured control NSC cultures after 30 days of differentiation. C4: Vimentin and neurofilament heavy immunoreactivity of SGZ uninjured control NSC cultures after 30 days of differentiation (C3 white box). D1: GFAP, Beta-III-Tubulin, and Map2 immunoreactivity of SGZ TBI-activated NSC cultures after 30 days of differentiation. D2: GFAP, Beta-III-Tubulin, and Map2 immunoreactivity of SGZ TBI-activated NSC cultures after 30 days of differentiation (D1 white box). D3: Vimentin and neurofilament heavy immunoreactivity of SGZ TBI-activated NSC cultures after 30 days of differentiation. D4: Vimentin and neurofilament heavy immunoreactivity of SGZ TBI-activated NSC cultures after 30 days of differentiation (D3 white box). Scalebars: 1&3: 200 µm, 2&4 50 µm.

NSCs from the SGZ TBI-activated NSC cultures settled and expanded, forming small colony clusters and single cell grouping in a similar manner to that of the SGZ uninjured control NSC cultures before differentiation (23 DIV, 0 dDiff). However, by 15 dDiff the SGZ TBI-activated NSC cultures had drastically expanded, going from approximately 10% confluence to 85-95% confluence. A large proportion of the cells retained NSC-like morphology, with debris from dead cells covering large parts of the cultures. Small pockets of cells adopting neuronal morphology were spread sporadically throughout the cultures (Fig. 4D2). By 30 dDiff the majority of these neuronal pockets had disappeared, resulting in 90-95% confluence of apparent cell debris (Fig. 4D3). Immunostaining of the cell cultures however revealed that the cell population consisted of a large number of vimentin-positive and GFAP-negative cells spread out in a large monolayer (Fig. 5D3-4), with smaller clusters of GFAP-positive glial cells spread sporadically throughout. Few Beta-III-Tubulin and Map2 positive neuronal cells were present throughout the cultures (Fig. 5D1-2), with a complete absence of more mature neurofilament-heavy neuronal cells (Fig. 5D3-4).

## Discussion

Both SVZ uninjured control and TBI-activated NSC cultures exhibited a similar initial settlement pattern, forming monolayer colonies with occasional multilayered clusters during expansion (Fig. 4A&B). However, during the differentiation process, significant differences emerged. SVZ uninjured control cultures maintained a stable and interconnected organization, with cells adopting more distinct neuronal and glial morphology over time (Fig. 4A). Immunolabelling after 30 days of differentiation revealed dense clusters of Beta-III-Tubulin and Map2-positive cells containing few, singular GFAP-positive glial cells situated within colonies of vimentin-positive cells with fibroblast-like morphology. Such fibroblast-like cells have previously been reported to thrive under NSC culturing conditions, displaying both NSC-specific behaviors and markers (Park et al 2012). In our study, such vimentin positive clusters appear to be associated with the neuronal colonies, potentially playing a supportive role during the differentiation of the SVZ-derived NSCs (Fig. 5A). Seemingly singular neurofilament heavy (NFH) positive neuronal cells extended long (<200 µm) processes between the vimentin-positive cell clusters, with the processes terminating in complex structures, potentially synapsing onto neuronal cells within the dense Beta-III-Tubulin and Map2-positive neuronal clusters (Fig. 5A).

In contrast, TBI-activated NSC cultures developed into large monolayer colonies, with less defined morphological profiles than the uninjured control cultures (Fig. 4B). Immunolabelling revealed that these monolayer cultures primarily consisted of vimentin-positive fibroblast-like cells, though without clustering, but rather with individual Beta-III-Tubulin and Map2-positive, and GFAP-positive neuronal and glial cells respectively, scattered throughout the culture (Fig. 5). Similar to the uninjured control NSC cultures, the TBI-activated cultures also contained seemingly singular NFH-positive neuronal cells also extending long (<200 µm) processes between the fibroblast-like colonies (Fig. 5A&B3&4). While the TBI-activated SVZ-derived NFH-positive neuronal cells also terminated their external processes within the fibroblast-like vimentin-positive colonies, these terminal structures displayed far less complexity than the corresponding cells observed in the uninjured control cultures (5A&B3&4). This observation suggests a reduced number of target neuronal cells in the colonies for synapse formation by NFH-positive cells in the TBI-activated culture colonies, as compared to the uninjured control. This observation is further corroborated by the absence of dense clusters of Beta-III-Tubulin and Map2-positive neurons in the TBI-activated NSC cultures (Fig. 5B1&2). Such a pattern may reflect a TBI-induced alteration in NSC behavior, potentially stemming from lesion-induced cell type-specific changes in gene expression and lineage trajectory within the SVZ-derived NSC population. This aligns with prior findings indicating the unique response of the SVZ neurogenic niche to traumatic brain injury (Chen et al 2021, Yoshiya et al 2003).

Similar to the SVZ, both SGZ uninjured control and TBI-activated conditions initially formed free-floating neurospheres that quickly adhered to the culture vessel surface. Studies into the function of the SGZ neurogenic niche have observed that the rate of proliferation and the fate of NSC progeny in the SGZ are strictly regulated by external factors such as vasculature and neuronal signaling (Bao et al 2017, Licht & Keshet 2015, Lin et al 2018). The SGZ receives modulating input from the medial septum that enhances NSC proliferative activity through GABAergic signaling, and ablation of this pathway results in a temporary increase of SGZ NSC proliferative activity and neurogenesis, which is followed by depletion of the SGZ NSC pool and impaired neurogenesis (Bao et al 2017, Káradóttir & Kuo 2018, Van der Borght et al 2005). We therefore thought it likely to observe a similar outcome of initial rapid NSC expansion followed by stable colony size, spontaneous differentiation and ultimately a decline in cell number due to the increased fragility of differentiating cells when passaging (Schiro et al 2022). However, we observed a drastic increase in cell growth from P1 to P2, which improved further by increasing the concentration of EGF in the expansion media.

The addition of EGF to the NSC expansion media was critical for survival and proliferation across both neurogenic niches. Homeostatic NSC-driven neurogenesis, particularly from the SVZ is mediated through an intermediary stage of DIx2-positive transit-amplifying C-type progenitor cells. EGF has previously been reported to convert these transit-amplifying C-type cells back to more NSC-like characteristics (Doetsch et al 1999, Doetsch et al 2002). We did not observe any DIx2 positive cells in either the SVZ or SGZ-derived uninjured control cultures at 20 DIV (S6). This potentially indicates either that NSC proliferation and neurogenesis may originate from other cell types, i.e., not requiring an intermediary stage or proliferation through other progeny cell types, or that conversion of transit-amplifying C-type cells back towards neural lineage multipotent NSCs takes place in the presence of EGF, as previously reported by Doetsch and colleagues (Doetsch et al 1999, Doetsch et al 2002). If the latter, then our findings support previous observations and indicate that this EGF-driven conversion of transit-amplifying C-type cells may provide a starting point for pharmacological intervention following TBI.

While differentiating, the uninjured control SGZ-derived NSCs initially formed small, adherent cell clusters that followed a similar developmental pattern to the SVZ-derived uninjured control cultures, but required longer time before organizing into semi-structured, interconnected colonies consisting of cells adopting distinct glial and neuronal morphology (Fig. 4C). Immunolabelling revealed these clusters to primarily consist of large vimentin-positive, but GFAP, Beta-III-Tubulin and Map2-negative cells adopting a fibroblast-like morphology with small clusters of GFAP-positive glial cells scattered throughout the cultures. Few, individual Beta-III-Tubulin and Map2-positive cells of neuronal morphology resided within the GFAP-positive clusters, extending processes between the clusters. While the SGZ-derived uninjured control cultures contained large vimentin-positive fibroblast-like cell clusters, similar to both SVZ-derived conditions, no neurofilament-heavy-positive neuronal cells were observed within or between the clusters observed in the SGZ-derived uninjured control NSC cultures.

The TBI-activated SGZ NSC cultures, however, exhibited a dramatic expansion, reaching high confluence after only 15 days of differentiation. However, unlike all the other conditions, large amounts of cell debris were present only within the TBI-activated cultures throughout the differentiation process (Fig. 4D1-3). Interestingly, this increased cell death restricted to the SGZ TBI-activated cultures only, resembles an increase in NSC progeny necrosis previously reported to occur within the SGZ following cortical impactor-induced TBI *in vivo* (Rola et al 2006, Ryu et al 2016, Zhou et al 2012). The cultures showed a predominance of cells with NSC-like morphology, and a notable absence of mature neuronal cells (Fig. 4D), suggesting that the early post-injury behavior of the NSCs within the SGZ may persist *in vitro* in some form beyond the harvesting and expansion phase. These behaviors may persist, and later become manifest during asymmetrical division when differentiation is no longer inhibited by the growth factors within the expansion media, given that this behavior was absent at all times in our SVZ and SGZ uninjured control cultures and SVZ TBI lesion-activated NSC cultures (Fig. 5A-D1-3).

While the addition of EGF may compensate for the GABAergic regulation of NSC proliferation, the loss of input from this pathway appears to result in restricted neurogenesis *in vitro*. Therefore, severing of this pathway, as occurs during the harvesting process of SGZ-derived NSCs may in part explain the observed rate of neural differentiation across both uninjured control and TBI-activated SGZ cultures compared to their SVZ-derived counterparts (Fig. 2&5, Table 1-2). Addition of VEGF to the cell culture media might to some extent improve NSC survival and neurogenesis beyond what was observed in this study (Fig. 5C&D (During & Cao 2006, Han et al 2015, Schänzer et al 2004, Segi-Nishida et al 2008)). Previous studies regarding the rate of SGZ-mediated neurogenesis in response to TBI indicated an increase in neurogenesis correlating with increased severity when TBI was induced through controlled cortical impact (Wang et al 2016). However, our results suggest this may not be the case for TBI lesions that directly impact the niche itself, with severing of the medal septum GABAergic projections to the SGZ likely to play a major role mediating the different responses. Interestingly, we found the SVZ-derived NSC cultures to still be highly neurogenic, even in the absence of the tight regulation of neurogenic activity *in vivo* (Xavier et al 2015), indicating potential differences at the cellular level as proposed by Petrik, D. et al (Petrik et al 2022).

Fibroblast-like cells have previously been reported to eventually outcompete NSCs in human-derived SVZ and SGZ cultures when cultured together, however, this has been reported to occur only after the 5^th^ or 6^th^ passage during expansion (Park et al 2012). In cultures derived from both neurogenic niches, these vimentin-positive cell types were observed. However, the neural cell immunoreactivity profile at the conclusion of differentiation varied considerably between the niches and experimental conditions (Fig. 5). This variation implies a potential disruption induced by the lesion, affecting both the fibroblast-like cells themselves and their potential interactions with NSCs. In the SVZ-derived uninjured control cultures, these cells appear to support dense Beta-III-Tubulin and Map2-positive neuronal colonies, a phenomenon not observed in the TBI-activated SVZ-derived cultures (Fig. 5A&B). This discrepancy may indicate altered cellular dynamics and intercellular communication within the SVZ following TBI.

The presence of fibroblast-like cells alone is however not sufficient to explain the high volume of cell death occurring solely in the SGZ-derived TBI-activated cultures during differentiation. As the fibroblast-like cells have been reported to be of a neurovascular origin (Park et al 2012), and they also display altered behavior in response to TBI. Our results may therefore indicate that cells involved in niche vasculature may also be affected by TBI, and further investigation into NSC-vasculature interactions following TBI may provide valuable insights into the potential of recruiting endogenous NSCs for neural tissue repair and replacement following TBI.

In our study, TBI was induced through a penetrative injury directly impacting the SGZ and the lesion did not increase neuronal differentiation, but influenced NSC differentiation toward a glial fate, resembling NSC behavior in response to TBI *in vivo* (Rola et al 2006). This suggests that the SGZ neurogenic niche either responds differently to direct insult, similar to the CC of the spinal cord niche (McDonough & Martínez-Cerdeño 2012, Meletis et al 2008), or neuronal cells are particularly susceptible to the necrotic environment induced by both within the niche (Rola et al 2006, Ryu et al 2016, Zhou et al 2012) and necrotic cues potentially still present in our *in vitro* cultures after TBI (Fig. 4D1-3). On the other hand, we cannot exclude that this may be an artifact of our protocol, as conditions were made to be as close as possible between the niches during the *in vitro* culturing process, thus warranting further investigation. This study underscores the complexity of NSC responses to TBI, revealing distinct outcomes in different neurogenic niches. These findings contribute to our understanding of how TBI affects neural lineage differentiation and the potential subsequent self-organization of NSC-derived progenitors, potentially informing future therapeutic strategies to promote recovery after brain trauma. However, further research is needed to elucidate the molecular mechanisms underlying these observed alterations in NSC behavior and differentiation patterns.

## Conclusion

Distinct proliferative and differentiation patterns arise between the uninjured control and TBI-activated NSCs from the two canonical neurogenic niches of the brain, primarily manifesting as increased proliferative potential among the TBI-activated NSCs. Such differences in the proportions of cells expressing Nestin, GFAP, Beta-III-Tubulin, and Map2, were also observed during both expansion and after 30 days of differentiation, indicating TBI-related changes in neural lineage commitment in NSCs from both the SVZ and SGZ. Our findings also highlight the impact of TBI on the potential for spontaneous organization of NSC progeny *in vitro*, with distinct profiles observed in cultures derived from uninjured control and TBI-activated NSCs from both neurogenic niches, thus providing valuable insights into the NSC responses to the early stage of traumatic brain injury.

## Author contributions

LES conceived the study, designed, and performed the experiments, collected, and analyzed the data, wrote, and finalized the manuscript; USB performed experiments, collected data, read, and edited the manuscript, CB performed brain sectioning, immunohistochemistry, Cresyl violet, and relevant imaging, read, and edited the manuscript; AS provided funding and guidance, revised, and edited the manuscript; IS provided funding and guidance, revised, and edited the manuscript.

## Acknowledgments

The authors would like to acknowledge support by the Liaison Committee for Education, Research, and Innovation in Central Norway (Samarbeidsorganet HMN-NTNU), the Joint Research Committee between St. Olav’s Hospital and the Faculty of Medicine and Health Sciences (Felles Forskningsutvalg), NTNU, and Enabling Technologies, NTNU. We thank Marit Trones Rem for technical assistance.

## Supplementary

**S1:** List of primary and secondary antibodies used.

**S1: List of primary and secondary antibodies used.**
| Antibody | Host species | Concentration | Identifier |
| --- | --- | --- | --- |
| Ki67 | Rabbit | 1:500 | ab15580 |
| Nestin | Mouse | 1:500 | ab6142 |
| Iba1 | Mouse | 1:500 | ab15690 |
| GFAP | Rabbit | 1:1000 | ab7260 |
| Nestin | Mouse | 1:500 | ab6142 |
| Ki67 | Rabbit | 1:500 | ab15580 |
| Map2 | Chicken | 1:20000 | ab5392 |
| Beta-III-Tubulin | Mouse | 1:1000 | ab78078 |
| GFAP | Rabbit | 1:1000 | ab7260 |
| FoxJ-1 | Mouse | 1:100 | SC-53139 |
| Host species | Wavelength | Concentration | Identifier |
| Anti-Rabbit | 488 | 1:500 | ab150077 |
| Anti-Mouse | 568 | 1:500 | ab175473 |
| Anti-Chicken | 647 | 1:500 | ab150171 |
| Hoechst 33342 | 461 | 1:2000 | C10337 G |

**Table S2:** NSC expansion over 20 DIV.

| Niche | Generation | Average number of live cells | St. Dev |
| --- | --- | --- | --- |
| SVZ Control | P0 | 500 000 | 0 |
|  | P1 | 387 000 | 74 953 |
|  | P2 | 5 120 000 | 113 137 |
| SVZ TBI | P0 | 500 000 | 0 |
|  | P1 | 1 180 000 | 84 853 |
|  | P2 | 10 720 000 | 1 527 351 |
| SGZ Control | P0 | 500 000 | 0 |
|  | P1 | 1 167 750 | 233 324 |
|  | P2 | 14 300 000 | 1 840 290 |
| SGZ TBI | P0 | 500 000 | 0 |
|  | P1 | 2 862 500 | 432 233 |
|  | P2 | 28 575 000 | 13 703 132 |

**Table S3:** Average number of cells positive for proliferative, NSC, and neural fate markers.

| Niche | Hoechst | EdU | Ki67 | Nestin | GFAP | B3T | Map2 |
| --- | --- | --- | --- | --- | --- | --- | --- |
| SVZ Control | 923.8 | 292.3 | 23.4 | 845.9 | 1.3 | 95.9 | 45.4 |
| St. Dev | 454.5 | 68.2 | 9.4 | 236.6 | 1.4 | 36.2 | 15.2 |
| SVZ TBI | 127.1 | 114.6 | 20.3 | 68.3 | 5.6 | 32.0 | 6.0 |
| St. Dev | 84.0 | 29.3 | 12.4 | 38.3 | 1.8 | 8.1 | 3.4 |
| SGZ Control | 349.9 | 280.4 | 5.6 | 161.1 | 9.3 | 5.7 | 42.1 |
| St. Dev | 162.0 | 163.6 | 2.8 | 39.4 | 2.4 | 2.1 | 17.2 |
| SGZ TBI | 239.8 | 124.1 | 29.0 | 154.2 | 3.8 | 2.3 | 28.0 |
| St. Dev | 93.0 | 56.4 | 13.6 | 38.8 | 2.0 | 1.4 | 9.9 |

**S4: TBI lesioning of the SVZ and SGZ neurogenic niches for local NSC injury response activation.**

**S4:**
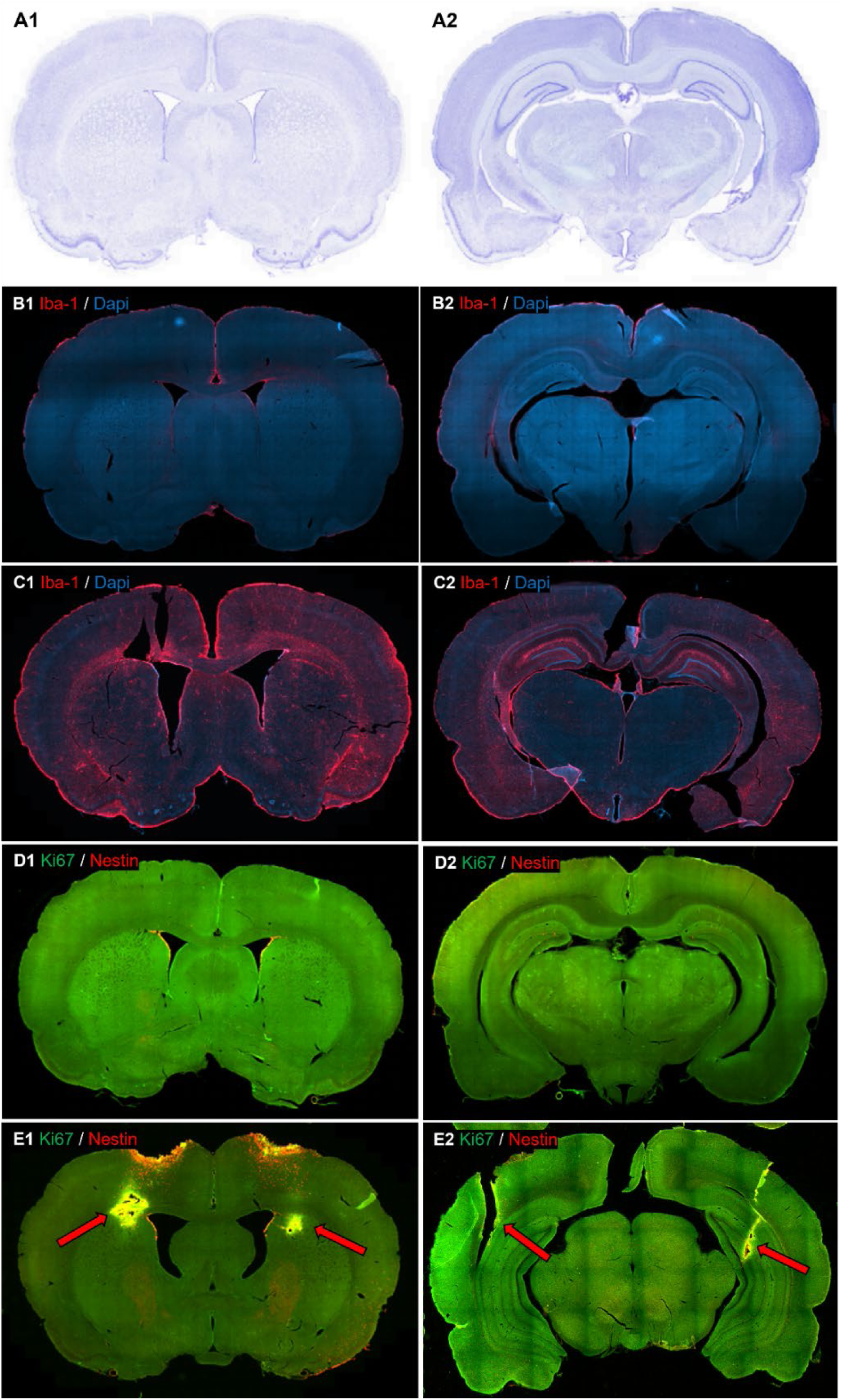
TBI lesioning of the SVZ and SGZ neurogenic niches for local NSC injury response activation. A: Nissl stain of uninjured control brain sections containing 1) SVZ and 2) SGZ neurogenic niches. B: Expression of reactive microglia (Iba-1; red) uninjured control SVZ (1) and SGZ (2) containing brain tissue. C: Expression of reactive microglia (Iba-1; red) in SVZ (1) and SGZ (2) TBI lesioned brain tissue. D: Baseline presence of active proliferation (Ki67; green) and Nestin-positive (red) NSCs in uninjured control SVZ (1) and SGZ (2) neurogenic niches. E: TBI lesion-dependent activation and migration of actively proliferating (Ki67; green) Nestin-positive NSCs towards lesion site (red arrows) in brains with TBI lesions targeting the SVZ (1) and SGZ (2) neurogenic niches.

**S5:**
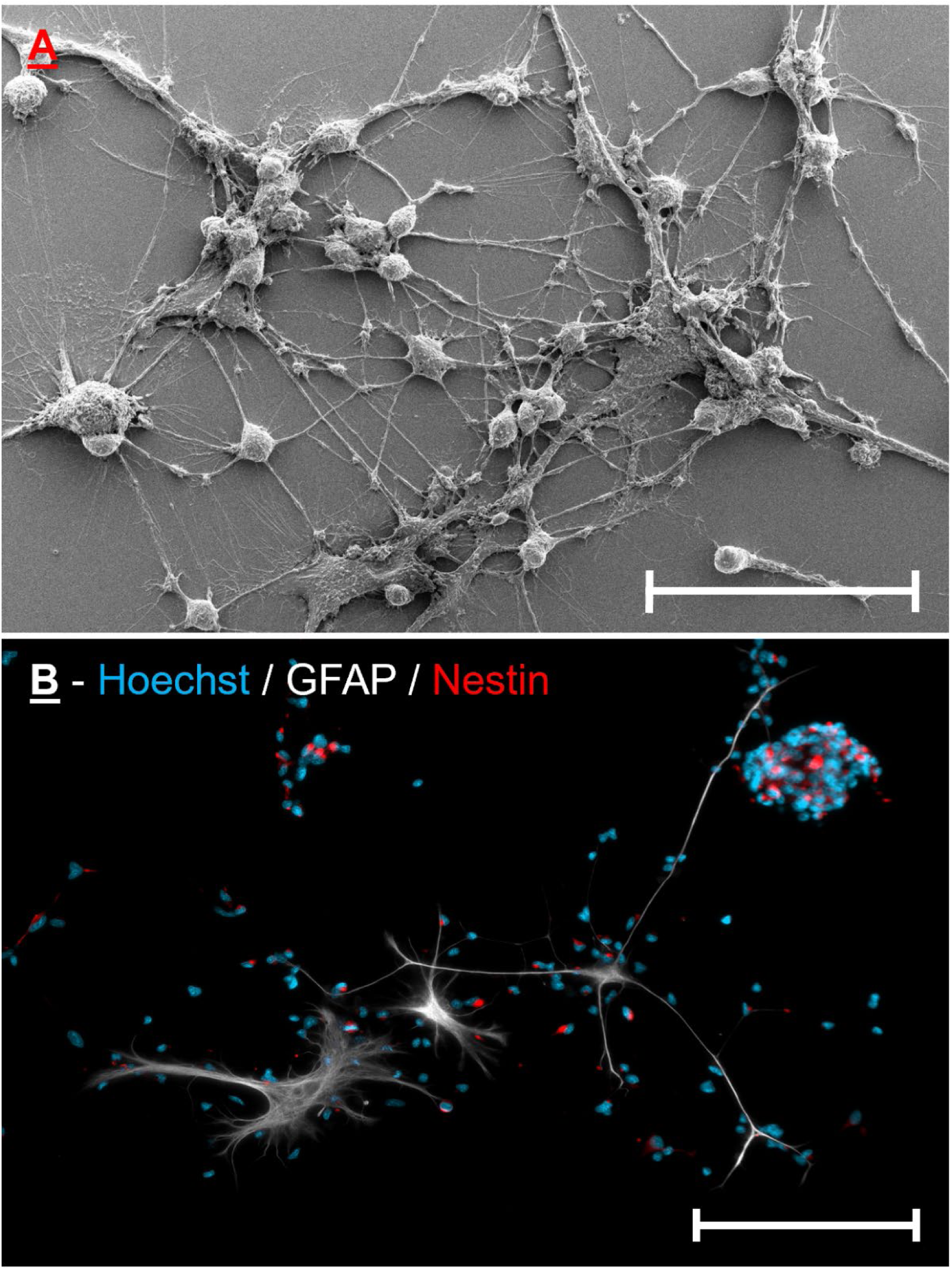
SGZ uninjured control NSCs settling along external processes of large GFAP positive cells. A: SEM image of NSCs settling around large GFAP-positive cells. Scalebar: 50 µm. B: Immunolabelling of NSCs settling around large GFAP-positive cells and in small clusters. Scalebar: 50 µm.

**S6.**
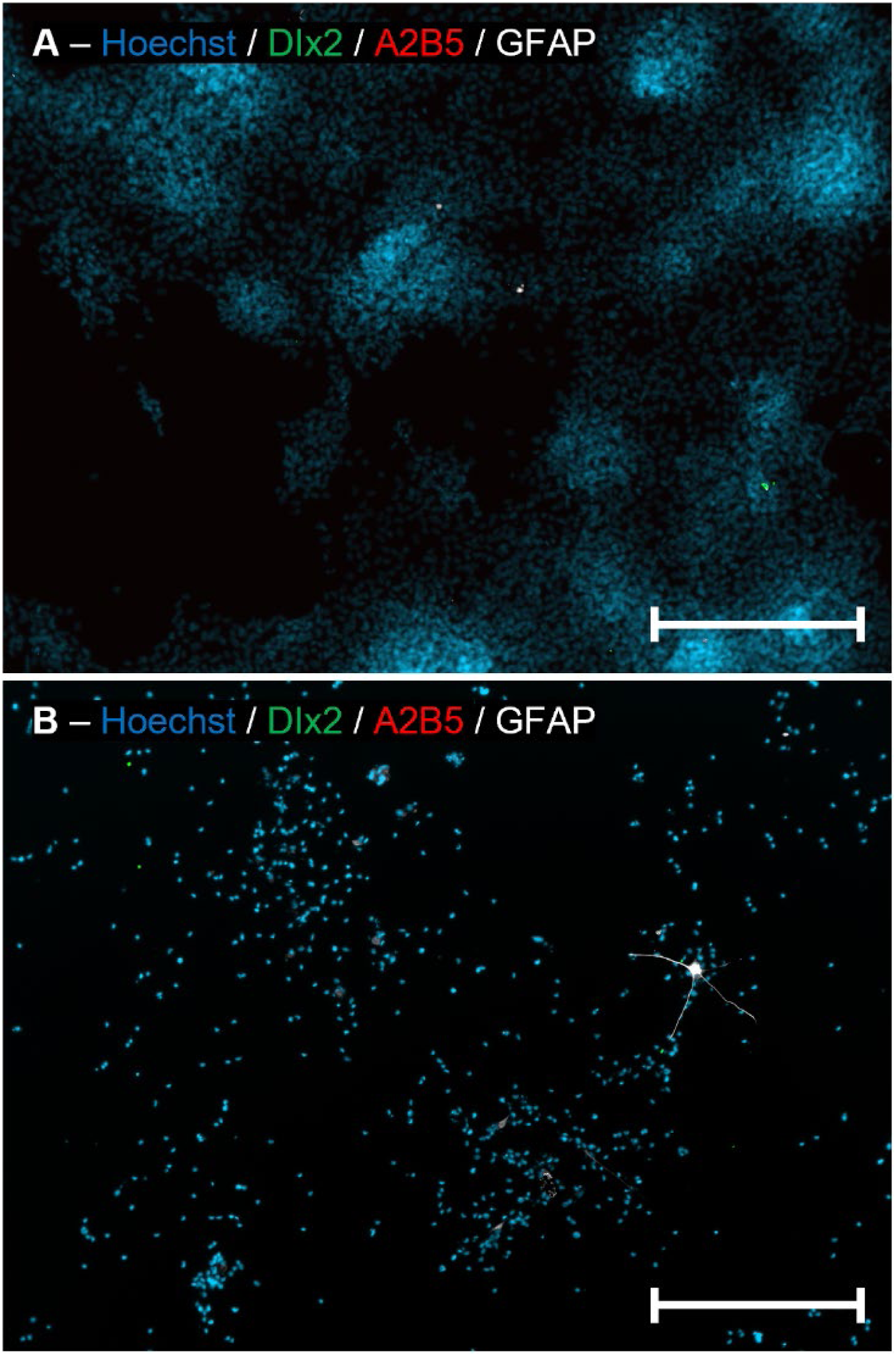
Absence of oligodendrocyte and transit amplifying precursor cells in SVZ and SGZ-derived NSC cultures. A: Immunolabelling of SVZ uninjured control NSC cultures indicating the absence of DIx2 positive transit-amplifying C-type cells and A2B5 positive oligodendrocyte progenitors. B: Immunolabelling of SGZ uninjured control NSC cultures indicating the absence of DIx2 positive transit-amplifying C-type cells and A2B5 positive oligodendrocyte progenitors. Scalebars: 200 µm.

## References

Abbott LC, Nigussie F. 2020. Adult neurogenesis in the mammalian dentate gyrus. Anatomia, histologia, embryologia 49: 3–16

Bao H, Asrican B, Li W, Gu B, Wen Z, et al. 2017. Long-range GABAergic inputs regulate neural stem cell quiescence and control adult hippocampal neurogenesis. Cell stem cell 21: 604–17. e5

Bazan E, Alonso F, Redondo C, Lopez-Toledano M, Alfaro JM, et al. 2004. In vitro and in vivo characterization of neural stem cells. Histology and Histopathology

Bjornsson Christopher S, Apostolopoulou M, Tian Y, Temple S. 2015. It Takes a Village: Constructing the Neurogenic Niche. Developmental Cell 32: 435–46

Chen X, Cao S, Wang Y, Li M, Guo Y, et al. 2021. Single-cell profiling resolved transcriptional alterations and lineage dynamics of subventricular zone after mild traumatic brain injury. bioRxiv: 2021.05. 31.446381

De Winter JC. 2019. Using the Student’s t-test with extremely small sample sizes. Practical Assessment, Research, and Evaluation 18: 10

Doetsch F. 2003. The glial identity of neural stem cells. Nature neuroscience 6: 1127–34

Doetsch F, Caillé I, Lim DA, García-Verdugo JM, Alvarez-Buylla A. 1999. Subventricular Zone Astrocytes Are Neural Stem Cells in the Adult Mammalian Brain. Cell 97: 703–16

Doetsch F, Petreanu L, Caille I, Garcia-Verdugo J-M, Alvarez-Buylla A. 2002. EGF Converts Transit-Amplifying Neurogenic Precursors in the Adult Brain into Multipotent Stem Cells. Neuron 36: 1021–34

During JM, Cao L. 2006. VEGF, a Mediator of the Effect of Experience on Hippocampal Neurogenesis. Current Alzheimer Research 3: 29–33

Gage FH. 2000. Mammalian neural stem cells. Science 287: 1433–38

Ghosh HS. 2019. Adult neurogenesis and the promise of adult neural stem cells. Journal of experimental neuroscience 13: 1179069519856876

Han J, Calvo C-F, Kang TH, Baker KL, Park J-H, et al. 2015. Vascular endothelial growth factor receptor 3 controls neural stem cell activation in mice and humans. Cell reports 10: 1158–72

Itoh T, Satou T, Hashimoto S, Ito H. 2005. Isolation of neural stem cells from damaged rat cerebral cortex after traumatic brain injury. Neuroreport 16: 1687–91

Itoh T, Satou T, Nishida S, Hashimoto S, Ito H. 2007. Immature and mature neurons coexist among glial scars after rat traumatic brain injury. Neurological research 29: 734–42

Johansson CB, Momma S, Clarke DL, Risling M, Lendahl U, Frisen J. 1999. Identification of a neural stem cell in the adult mammalian central nervous system. Cell 96: 25–34

Káradóttir RT, Kuo CT. 2018. Neuronal activity-dependent control of postnatal neurogenesis and gliogenesis. Annual review of neuroscience 41: 139–61

Kennea NL, Mehmet H. 2002. Neural stem cells. The Journal of Pathology: A Journal of the Pathological Society of Great Britain and Ireland 197: 536–50

Licht T, Keshet E. 2015. The vascular niche in adult neurogenesis. Mechanisms of Development 138: 56–62

Lim DA, Alvarez-Buylla A. 2016. The adult ventricular–subventricular zone (V-SVZ) and olfactory bulb (OB) neurogenesis. Cold Spring Harbor perspectives in biology 8: a018820

Lin R, Cai J, Kenyon L, Iozzo R, Rosenwasser R, Iacovitti L. 2018. Systemic Factors Trigger Vasculature Cells to Drive Notch Signaling and Neurogenesis in Neural Stem Cells in the Adult Brain. Stem Cells 37: 395–406

McDonough A, Martínez-Cerdeño V. 2012. Endogenous proliferation after spinal cord injury in animal models. Stem cells international 2012

Meletis K, Barnabé-Heider F, Carlén M, Evergren E, Tomilin N, et al. 2008. Spinal cord injury reveals multilineage differentiation of ependymal cells. PLoS biology 6: e182

Mizrak D, Levitin HM, Delgado AC, Crotet V, Yuan J, et al. 2019. Single-cell analysis of regional differences in adult V-SVZ neural stem cell lineages. Cell reports 26: 394–406. e5

Obernier K, Alvarez-Buylla A. 2019. Neural stem cells: origin, heterogeneity and regulation in the adult mammalian brain. Development 146: dev156059

Park TI-H, Monzo H, Mee EW, Bergin PS, Teoh HH, et al. 2012. Adult human brain neural progenitor cells (NPCs) and fibroblast-like cells have similar properties in vitro but only NPCs differentiate into neurons. PloS one 7: e37742

Paxinos G, Watson C. 1997. Paxinos and Watson’s the rat brain in stereotaxic coordinates. (No Title)

Petrik D, Jörgensen S, Eftychidis V, Siebzehnrubl FA. 2022. Singular Adult Neural Stem Cells Do Not Exist. Cells 11: 722

Price J, Williams BP. 2001. Neural stem cells. Current Opinion in Neurobiology 11: 564–67

Redmond SA, Figueres-Oñate M, Obernier K, Nascimento MA, Parraguez JI, et al. 2019. Development of ependymal and postnatal neural stem cells and their origin from a common embryonic progenitor. Cell reports 27: 429–41. e3

Richter KN, Revelo NH, Seitz KJ, Helm MS, Sarkar D, et al. 2018. Glyoxal as an alternative fixative to formaldehyde in immunostaining and super-resolution microscopy. The EMBO journal 37: 139–59

Rola R, Mizumatsu S, Otsuka S, Morhardt DR, Noble-Haeusslein LJ, et al. 2006. Alterations in hippocampal neurogenesis following traumatic brain injury in mice. Experimental neurology 202: 189–99

Ryu JR, Hong CJ, Kim JY, Kim E-K, Sun W, Yu S-W. 2016. Control of adult neurogenesis by programmed cell death in the mammalian brain. Molecular Brain 9: 43

Salman H, Ghosh P, Kernie SG. 2004. Subventricular zone neural stem cells remodel the brain following traumatic injury in adult mice. Journal of neurotrauma 21: 283–92

Schänzer A, Wachs F-P, Wilhelm D, Acker T, Cooper-Kuhn C, et al. 2004. Direct Stimulation of Adult Neural Stem Cells In Vitro and Neurogenesis In Vivo by Vascular Endothelial Growth Factor. Brain Pathology 14: 237–48

Schiro LE, Bauer US, Sandvig A, Sandvig I. 2022. Isolation and comparison of neural stem cells from the adult rat brain and spinal cord canonical neurogenic niches. STAR Protocols 3: 101426

Segi-Nishida E, Warner-Schmidt JL, Duman RS. 2008. Electroconvulsive seizure and VEGF increase the proliferation of neural stem-like cells in rat hippocampus. Proceedings of the National Academy of Sciences 105: 11352

Shihabuddin LS, Horner PJ, Ray J, Gage FH. 2000. Adult spinal cord stem cells generate neurons after transplantation in the adult dentate gyrus. Journal of Neuroscience 20: 8727–35

Snedecor GW, Cochran WG. 1989. Statistical Methods, eight edition. Iowa state University press, Ames, Iowa 1191

Song H-j, Stevens CF, Gage FH. 2002. Neural stem cells from adult hippocampus develop essential properties of functional CNS neurons. Nature neuroscience 5: 438–45

Sotocina Susana G, Sorge Robert E. 2011. Zaloum Austin, Tuttle Alexander H, Martin Loren J, Wieskopf Jeffrey S, Mapplebeck Josiane CS, Wei Peng, Zhan Shu, Zhang Shuren, McDougall Jason J, King Oliver D, Mogil Jeffrey S. The Rat Grimace Scale: A Partially Automated Method for Quantifying Pain in the Laboratory Rat via Facial Expressions. Molecular Pain 7: 1744–8069

Taupin P. 2006. Adult neural stem cells, neurogenic niches, and cellular therapy. Stem Cell Reviews 2: 213–19

Van der Borght K, Mulder J, Keijser JN, Eggen BJ, Luiten PG, Van der Zee EA. 2005. Input from the medial septum regulates adult hippocampal neurogenesis. Brain research bulletin 67: 117–25

Wang X, Gao X, Michalski S, Zhao S, Chen J. 2016. Traumatic brain injury severity affects neurogenesis in adult mouse hippocampus. Journal of neurotrauma 33: 721–33

Xavier ALR, Kress BT, Goldman SA, de Menezes JRL, Nedergaard M. 2015. A distinct population of microglia supports adult neurogenesis in the subventricular zone. Journal of Neuroscience 35: 11848–61

Yoshiya K, Tanaka H, Kasai K, Irisawa T, Shiozaki T, Sugimoto H. 2003. Profile of gene expression in the subventricular zone after traumatic brain injury. Journal of neurotrauma 20: 1147–62

Young SZ, Taylor MM, Bordey A. 2011. Neurotransmitters couple brain activity to subventricular zone neurogenesis. European Journal of Neuroscience 33: 1123–32

Zhou H, Chen L, Gao X, Luo B, Chen J. 2012. Moderate Traumatic Brain Injury Triggers Rapid Necrotic Death of Immature Neurons in the Hippocampus. Journal of Neuropathology & Experimental Neurology 71: 348–59

